# Calcium stabilization of a flexible N-terminal domain in a pentameric ligand-gated ion channel

**DOI:** 10.1101/2025.02.24.640012

**Authors:** Chen Fan, Marie Lycksell, Yuxuan Zhuang, Rebecca J. Howard, Erik Lindahl

**Affiliations:** Dept. of Applied Physics, Science for Life Laboratory, KTH Royal Institute of Technology, Solna, Sweden; Dept. of Biochemistry and Biophysics, Science for Life Laboratory, Stockholm University, Solna, Sweden; Department of Pharmacology and Chemical Biology, Shanghai Jiao Tong University School of Medicine, Shanghai, China

**Keywords:** ligand-gated ion channel, cryo-EM, molecular dynamics, calcium

## Abstract

Pentameric ligand-gated ion channels (pLGICs) are responsible for the rapid conversion of chemical to electrical signals. In addition to the canonical extracellular and transmembrane domains, some prokaryotic pLGICs contain an N-terminal domain (NTD) of unclear structure and function. In one such case, the calcium-sensitive channel DeCLIC, the NTD appears to accelerate gating; however, its evident flexibility has posed a challenge to model building, and its role in calcium sensitivity is unclear. Here we report cryo-EM structures of DeCLIC to the highest resolutions thus far, in circularized lipid nanodiscs. Application of refinement tools enabled definition of multiple calcium-binding sites in each symmetric subunit. Under calcium-free conditions, both symmetric and asymmetric classes could be reconstructed, implicating a heterogeneous ensemble. Behavior of these structures in molecular dynamics simulations was consistent with calcium stabilization of the NTD, an effect that may be conserved in structurally homologous domains across evolution.

## Introduction

Pentameric ligand-gated ion channels (pLGICs) constitute a family of receptors responsible for the rapid conversion of chemical to electrical signals in most kingdoms of life.^1^ In animals, pLGICs correspond to the historical Cys-loop receptor family, whose members specifically recognize the neurotransmitters acetylcholine, ɣ-aminobutyric acid, serotonin, or glycine.^2^ Consistent with their physiological importance, these ion channels are important pharmaceutical targets in the treatment of psychiatric disorders including epilepsy, depression, anxiety, and autism.^3^

Based on sequence alignment with eukaryotic Cys-loop receptors, pLGICs have also been identified in a range of prokaryotes. These homologs show evidence of conserved topologies and gating mechanisms, and may offer straightforward model systems to study structure-function relationships. Indeed, the first full-length X-ray structures reported in this family were of the bacterial channels ELIC and GLIC.^4–6^ These structures confirm an overall fivefold (pseudo)symmetric assembly, each subunit containing ten β-strands (β1–β10) that contribute to an extracellular or periplasmic domain (ECD), and four helices (M1–M4) that contribute to a transmembrane domain (TMD) (Fig. S1). The M2 helices line the ion-conducting transmembrane pore, typically constricted by pore-facing amino-acid side chains at the so-called 2’, 9’, and 16’ positions. Upon activation by ligands or protons, a series of allosteric conformational changes including relative rotation of the ECD (ECD twist) and remodeling of domain-interface β-strands (β-expansion) eventually opens a pore in the TMD.^1^ Compared with their eukaryotic counterparts, prokaryotic pLGICs typically lack a substantial intracellular domain (ICD), but may include an N-terminal domain (NTD) localized to the periplasmic side of the membrane, about which little is known.^7,8^

To date, the only pLGIC in which an NTD has been partially characterized is DeCLIC, derived from a symbiotic sulfate-reducing deltaproteobacterium related to *Desulfofustis glycolicus*. The DeCLIC NTD accounts for roughly half the total receptor mass (Fig. 1A, S1), and appears to accelerate activation and deactivation of the channel.^9^ Within each subunit, the NTD consists of two lobes (NTD1 and NTD2) each adopting a jelly-roll fold. In both X-ray and electron cryomicroscopy (cryo-EM) structures of five-fold symmetric DeCLIC in detergent,^9,10^ density in the NTD is poorly defined relative to the ECD and TMD, suggesting this domain is partially unstructured, flexible, and/or heterogeneous. Indeed, unrestrained molecular dynamics (MD) simulations spontaneously generate multiple asymmetric conformations of the NTD, largely via rigid-body rearrangements of individual NTD1 lobes away from the periplasmic vestibule structures.^10^ Moreover, solution-phase small-angle scattering measurements are more consistent with these MD-generated asymmetric conformations than with symmetric X-ray or cryo-EM structures.^10^

**Figure 1.**
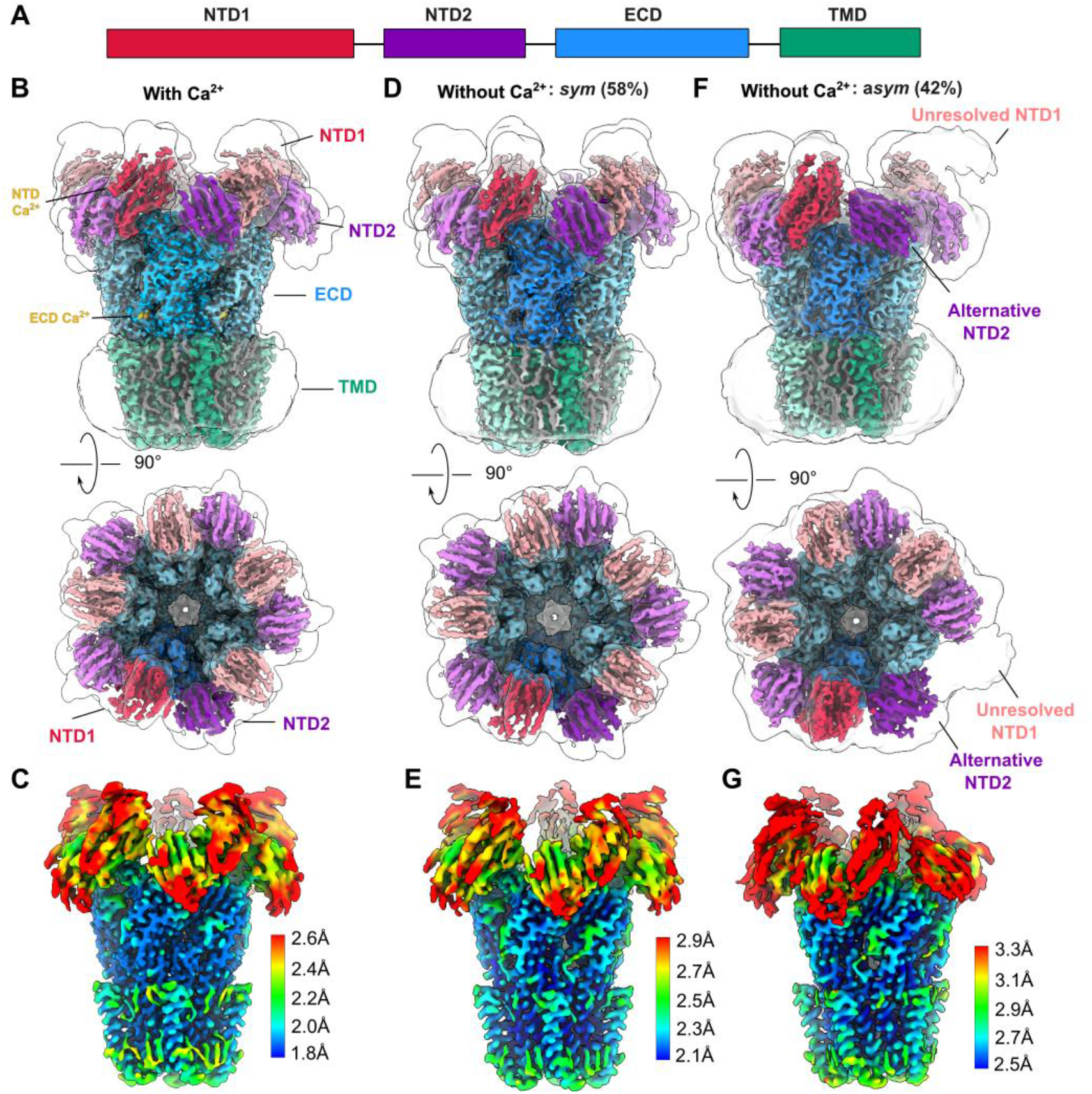
Calcium depletion promotes an asymmetric-NTD conformation of DeCLIC. A) Domain architecture of a single DeCLIC subunit gene, including NTD1 (red) and NTD2 (purple) lobes and contributions to the ECD (blue) and TMD (green). B) Cryo-EM map of DeCLIC in the presence of calcium, viewed from the membrane plane (above) and from the periplasmic side (below). Structural domains are labeled and colored as in *A*, along with non-protein densities modeled as lipids (gray) and Sites 1, 2, 3, and 4 (gold). The same map, low-pass filtered at 8 Å, is shown as a transparent envelope. C) Map as in *B*, viewed from the membrane plane and colored by local resolution according to the scale bar at right. D) Map of DeCLIC corresponding to the *sym* conformation in the absence of calcium, depicted as in *B*. E) Map as in *D*, depicted as in *C*. F) Map of DeCLIC corresponding to the *asym* conformation in the absence of calcium, depicted as in *B*. E) Map as in *F*, depicted as in *C*.

Like several pLGICs, DeCLIC has been shown to be inhibited by calcium ions (Ca^2+^), mediated at least in part by a calcium-binding site at the ECD-TMD interface.^9^ The NTD surface also contains several regions of negative charge, and includes sites of anomalous X-ray scattering consistent with calcium binding in at least some subunits^9^; however, the specific relationship between calcium and NTD dynamics, if any, remains largely undefined. Whereas DeCLIC conducts currents in response to calcium depletion in *Xenopus* oocytes,^9^ a potentially open X-ray structure determined in the absence of calcium was inconsistent with MD and small-angle scattering data, and all cryo-EM structures reported thus far have contained closed pores.^10^ Thus, open questions remain as to the experimental conditions required for channel gating.

Here we report cryo-EM structures of DeCLIC in the absence and presence of calcium in circularized nanodiscs, achieving higher overall resolution (2.1-2.2 Å) than previously reported (3.2-3.8 Å). As observed under similar conditions in detergent, the channel contains a closed pore, with weaker density in the NTD relative to other domains. Under calcium-chelating conditions, an additional class could be reconstructed in which one set of adjacent NTD1 and NTD2 lobes was even more poorly resolved and displaced from the symmetric pentamer, consistent with a heterogeneous asymmetric ensemble in the absence of calcium. Application of tools including 3D Flexible Refinement (3DFlex) in CryoSPARC facilitated model building and identification of multiple calcium-binding sites in the NTD and ECD of each symmetric subunit. Behavior of these structures in MD simulations was consistent with calcium binding stabilizing the NTD overall, an effect that may be conserved in structurally homologous domains across evolution.

## Results

### Calcium depletion promotes an asymmetric-NTD conformation of DeCLIC

In order to elucidate heterogeneity and possible calcium interactions of the DeCLIC NTD, we sought to improve the resolution of our cryo-EM structures by altering the membrane mimetic and applying recently developed refinement methods. Whereas in all previously reported structures, DeCLIC was solubilized in detergent,^9,10^ here we reconstituted the purified protein with soybean polar lipids. The lipid belt was stabilized by a circularized nanodisc with a predicted diameter of ∼15 nm,^11^ chosen to be larger than that predicted for the channel.

We first collected data in the presence of 10 mM calcium, conditions shown to inhibit channel function.^9^ From this dataset, a single well defined class could be refined with five-fold symmetry to an overall resolution of 2.1 Å (Fig. 1B, S2, S3, Table 1). As expected, the channel contained a closed pore (Fig. 1B, S1). Although local resolution in the NTD was poor relative to the ECD and TMD (Fig. 1C), all residues in the NTD1 and NTD2 lobes could be confidently assigned. Further application of 3DFlex^12^ without imposing symmetry dramatically improved the density, enabling clear definition of side chains as well as β-strand backbones for most residues (Fig. S4). Following application of the deep learning-based post-processing tool DeepEMhancer^13^, > 90% of NTD residues could be built *de novo*, including some modifications relative to previous structures in the loops linking the NTD and ECD (residues 98-105, 121-130, 153-161) (Fig. 3C). Aside from the NTD, the most apparent difference between this nanodisc structure and previous detergent structures was a modest contraction at the inner end of the TMD pore (2.5 Å to 1.5 Å radius at 2’, Fig. S1).

We then collected data in the absence of calcium, ensured by adding 10 mM ethylenediaminetetraacetic acid (EDTA) to the nanodisc-reconstituted sample. In order to identify potentially asymmetric conformations, we initially processed the resulting dataset without imposing symmetry, and identified two classes. Although both contained non-conducting pores similar to the calcium, one of these (*sym*, 2.4 Å, 58% of assigned particles) exhibited five-fold symmetry and was further refined in C5 (Fig. 1D, S1, S2, S3, Table 1). The NTD density, albeit weak, enabled assignment of all NTD1 and NTD2 residues (Fig. 1E). In a second class (*asym*, 2.7 Å, 42% of assigned particles), one NTD1 lobe was even more poorly resolved than the others, only clearly visible in a low-pass filtered map (Fig. 1F–G, S2, S3, Table 1). Still, application of 3DFlex and DeepEMhancer (Fig. S4) enabled building of > 90% of NTD residues in both conformations.

### Calcium-binding sites in the DeCLIC NTD and ECD

The electrostatic potential along the surface of DeCLIC is negatively charged in several regions of the NTD as well as ECD (Fig. S5), suggesting multiple possible interaction sites for Ca^2+^. A previous X-ray structure of DeCLIC derived from crystals soaked in 200 mM Ba^2+^ also includes sites of anomalous diffraction in NTD1, though detailed protein-ion interactions in this region have not been defined.^9^ Here, we observed four roughly spherical non-protein densities associated with each subunit, appearing in the presence but not the absence of calcium; we named these Sites 1, 2, 3, and 4 (Fig. 2A–C, S6). Given the limited resolution and substantial negative potential distributed across the DeCLIC NTD (Fig. 2C, S5A), these densities may not represent all prospective Ca^2+^ interactions, rather those that could be characterized with relatively high confidence.

**Figure 2.**
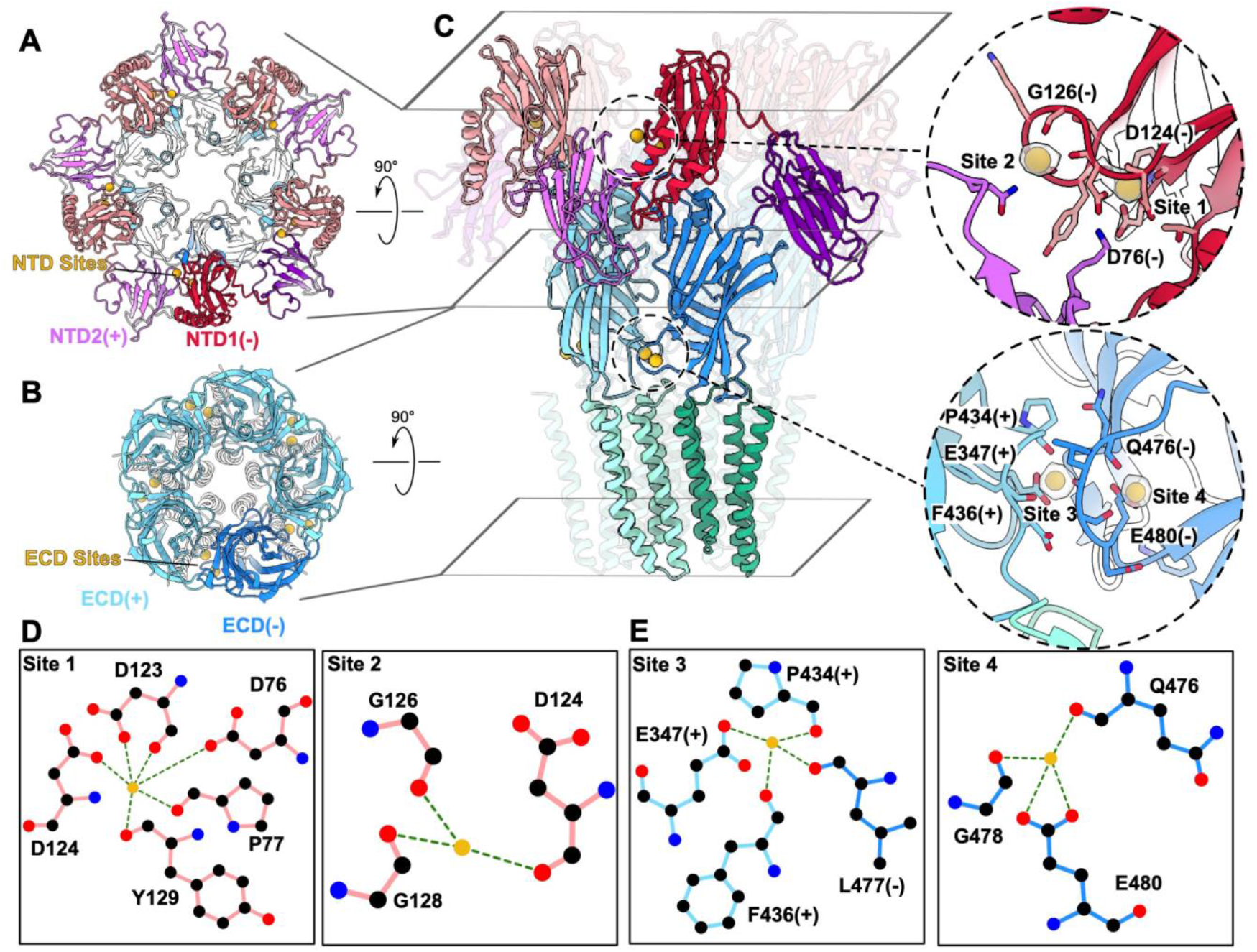
Prospective calcium-binding sites in the DeCLIC NTD and ECD. **A)** NTD and ECD of DeCLIC in the presence of calcium, viewed from the periplasmic side and colored as in *Fig. 1B*. For clarity, the lowermost subunit is shaded darker. Non-protein densities and proximal NTD lobes are labeled at the lowermost interface between principal (+) and complementary (−) subunits. **B)** ECD and TMD of DeCLIC in the presence of calcium, depicted as in *A*. **C)** Full-length DeCLIC in the presence of calcium, depicted as in *A–B* but viewed from the membrane plane, with the distal three subunits rendered transparent. Rhomboids indicate clipping planes used in *A–B*. Insets contain zoom views of non-protein densities associated with a single-subunit NTD (above) or ECD interface (below), as indicated in the full-length structure by dashed circles. **D)** Schematic of coordination of Sites 1 (left) and 2 (right) in a single NTD1 lobe. **E)** Schematic of coordination of Sites 3 (left) and 4 (right) at a single ECD interface.

**Figure 3.**
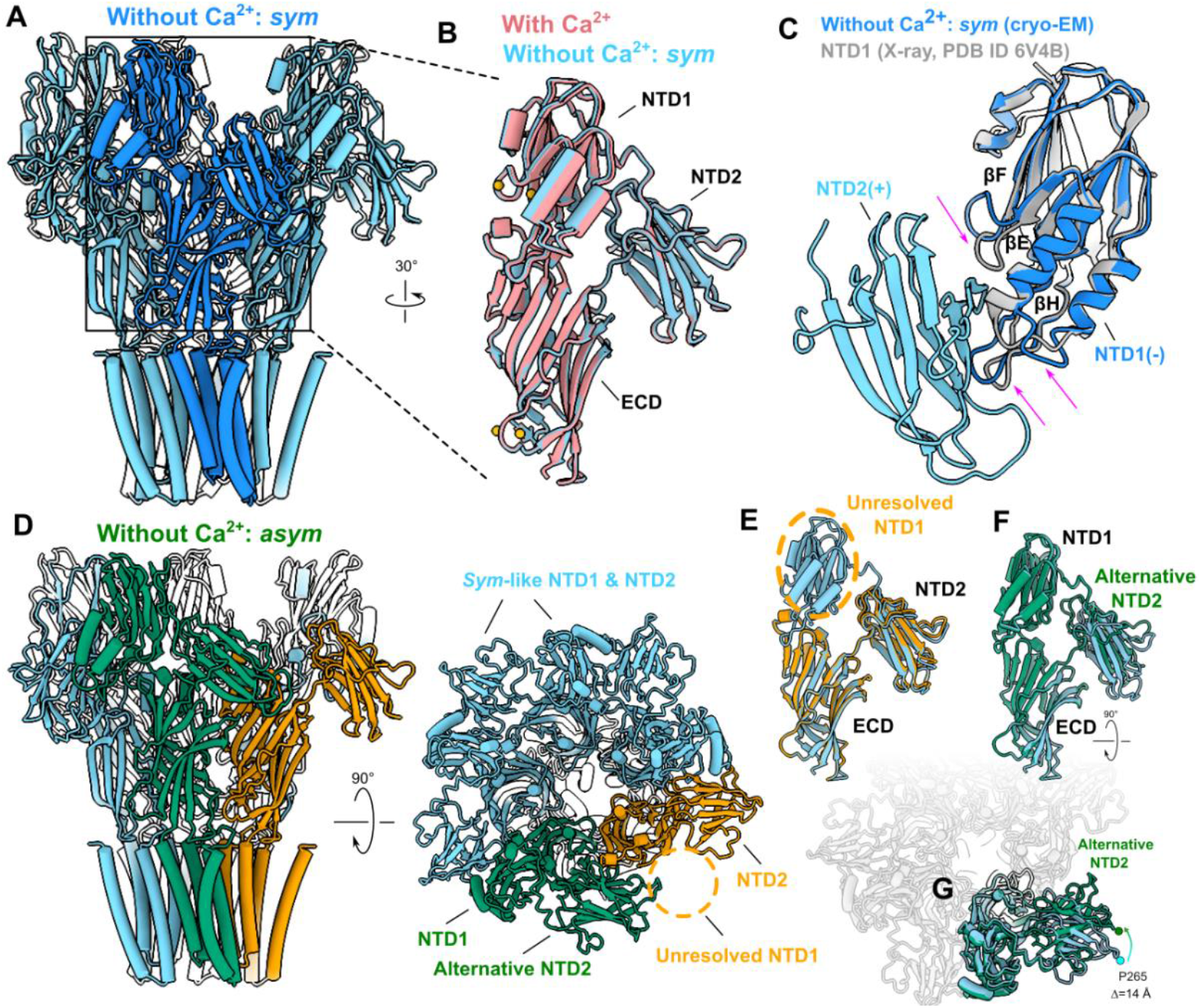
Remodeling of subunit interfaces surrounding calcium sites. **A)** DeCLIC structure correspon**ding to the *sym* conformation in the absence of calcium (blue)**, viewed from the membrane plane. For clarity, the foremost subunit is shaded darker. **B)** Superimposed structures of the NTD–ECD region of a single DeCLIC subunit with calcium (red) and in **the *sym* conformation without calcium** (blue). **C)** Structure of NTD1 determined as an isolated protein (gray) superimposed with the same subdomain in the context of the ***sym* DeCLIC conformation (blue). Magenta arrows indicate modest loop displacement in the isolated-lobe versus full-length structures**. NTD2 from the principal subunit (light blue) is included for perspective. **D)** DeCLIC structure correspon**ding to the *asym* conformation in the absence of calcium**, viewed from the membrane plane (left) and from the periplasmic side (right). Three subunits (blue) are comparable to the *sym* conformation; one (green) contains an alternative pose of NTD2; and in one (gold), NTD1 is too poorly defined for model building. **E)** NTD-ECD region in the *asym* subunit containing an unresolved NTD1 (gold), superimposed with the same region in **the *sym* conformation** (blue). **F)** NTD-ECD region in the *asym* subunit containing an alternative NTD2 pose (green), superimposed with the same region in **the *sym* conformation** (blue). **G)** Superposition as in *F* viewed from the periplasmic side, with neighboring subunits rendered partially transparent for perspective. To better show the translocation, P265 in each subunit is highlighted by beads and the distance is labeled.

As previously reported,^9^ each NTD lobe adopted a jelly-roll fold consisting of two four-stranded antiparallel β-sheets (strands βA, βH, βC, βF and βB, βG, βD, βE). NTD1 also contained two α-helices inserted in the βD–βE loop, and one α-helix at the C-terminus. Our present densities enabled building of all β-strands, α-helices, and intervening loops in both NTD lobes in each subunit, except the NTD2 βE-βF loop (residues 291-296) (Fig. 2A). Non-protein densities in each NTD subunit were proximal to the previously reported anomalous Ba^2+^ diffraction density,^9^ coordinated by the NTD1 βD, βE, and βF strands (Fig. 2C). These densities faced the periplasmic vestibule, occupying the complementary face of NTD1 proximal to NTD2 from the neighboring subunit (Fig. 2A, 2C, S5B, S6A). Two types of NTD densities were observed, one of which (Site 1) was coordinated by the acidic side chains of βD-Asp76, βE-Asp123, and βE-Asp124, as well as backbone carbonyls of βD-Pro77, βE-Asp123, and βE-Tyr129. A second density (Site 2) was more loosely coordinated by backbone carbonyls of βE-Asp124, βE-Gly126, and βF-Gly128 (Fig. 2C–D).

We also observed two non-protein densities at each ECD subunit interface. One (Site 3) was coordinated between the principal β1–β2 loop and complementary β8–β9 loop (historically called loop F), while the other (Site 4) was contained within loop F (Fig. 2B–C, S5C, S6B). Site 3 was coordinated by the acidic side chain of β1–β2 Glu347, as well as backbone carbonyls of β1–β2 Pro434, β1–β2 Phe436, and loop-F Leu477, and was similar to the Ca^2+^ site previously reported in lower-resolution detergent structures^9,10^. Site 4 was exposed to solvent and coordinated by the side chain of Glu480 and backbone carbonyls of Gln476 and Gly478, all in loop F (Fig. 2C, E).

### Remodeling of subunit interfaces around calcium sites

Our cryo-EM densities also enabled modeling of two classes of DeCLIC in the absence of calcium. Aside from the absence of ion-like densities in Sites 1, 2, 3, and 4, the *sym* calcium-free conformation was largely superimposable with the calcium-bound structure (Fig. 3A–B, S1, S6A–B). The most notable difference in the absence of calcium was a local loss of definition in the density associated with loop F (Fig. S6C), indicating calcium may stabilize this region in a closed-pore state. For reference, we also compared the *sym* conformation of NTD1 to a previously reported X-ray structure of the same subdomain as an isolated soluble protein (Fig. 3C).^9^ Secondary structure elements in these models were effectively identical, although relative compaction of the βE-βF and βG-βH loops in the isolated X-ray structure suggested that interactions with neighboring NTD2 or ECD regions influences local topology near the domain interface.

Aside from the absence of ion-like densities, the *asym* calcium-free conformation differed from the calcium structure at one NTD1 and adjacent NTD2 lobes (Fig. 1B, 2F, 3D–G). The asymmetric NTD2 lobe was associated with relatively poor density, but could be built in an alternative pose, displaced upward (away from the membrane plane) and rotated counter-clockwise (viewed from the periplasm) relative to the *sym*-like arrangement (Fig. 3F–G). The NTD1 lobe complementary to this asymmetric NTD2 exhibited even lower local resolution; although its poor definition precluded model building, low-pass filtering indicated a pose displaced outward from the central axis (Fig. 1F, 3D–E). Interestingly, a similar translocation was predicted by previous MD simulations and small-angle scattering measurements.^10^ Thus, depletion of calcium from the NTD2(+)/NTD1(−) interface enabled their partial mobilization and displacement in a subset of particles.

### Destabilization of closed DeCLIC upon calcium depletion

Given that calcium depletion appeared to promote an alternative conformation of DeCLIC, containing a pair of poorly resolved, potentially flexible NTD lobes, we next tested the effect of calcium on domain dynamics using MD simulations (Table 2). In quadruplicate >300-ns all-atom simulations of the DeCLIC structure with calcium, Ca^2+^ ions placed in both Sites 1 and 3 sites remained within 5 Å of their initial poses in at least four of five chains (Fig. S7). Ca^2+^ was less stable in Sites 2 and 4 sites, presumably reflecting their less extensive coordination (Fig. 2D-E, S7). Based on these observations, Sites 1 and 3 appear likely to be occupied by Ca^2+^ at relatively high affinity, while Sites 2 and 4 may be occupied at relatively low affinity and/or by alternative ligands.

Whereas no major changes were observed in the ECD-TMD regions in simulations with or without calcium (Fig. 4C-E), the NTD exhibited more flexibility, particularly upon calcium depletion (Fig. 4F-H). In simulations of the calcium-bound structure, the center-of-masses (COMs) of individual NTD1 lobes deviated up to 13 Å from their initial positions; in simulations of the structure without calcium (in the *sym* conformation), such deviations increased up to 25 Å (Fig. 4A, 4F, S8A). Although NTD1 lobes sampled multiple poses, in several frames they transitioned outward from the channel axis, consistent with low-resolution density in the *asym* cryo-EM structure (Fig. 1F). Similarly, NTD2 lobes deviated up to 7 Å with calcium, but up to 15 Å without calcium (Fig. 4B, 4G, S8B). Global properties also exhibited modest destabilization upon calcium removal: from an initial value of −26°, ECD twist varied in a 7° range with calcium, but in a 12° range without calcium (Fig. S8C). Similarly, β-expansion varied in a range of 4 Å versus 6 Å in the presence and absence of calcium, respectively (Fig. 4E, S8D). Thus, our structures and simulations indicate that calcium depletion destabilizes the DeCLIC closed state, as evidenced by global structural metrics associated with gating^14^ and particularly the NTD.

**Figure 4.**
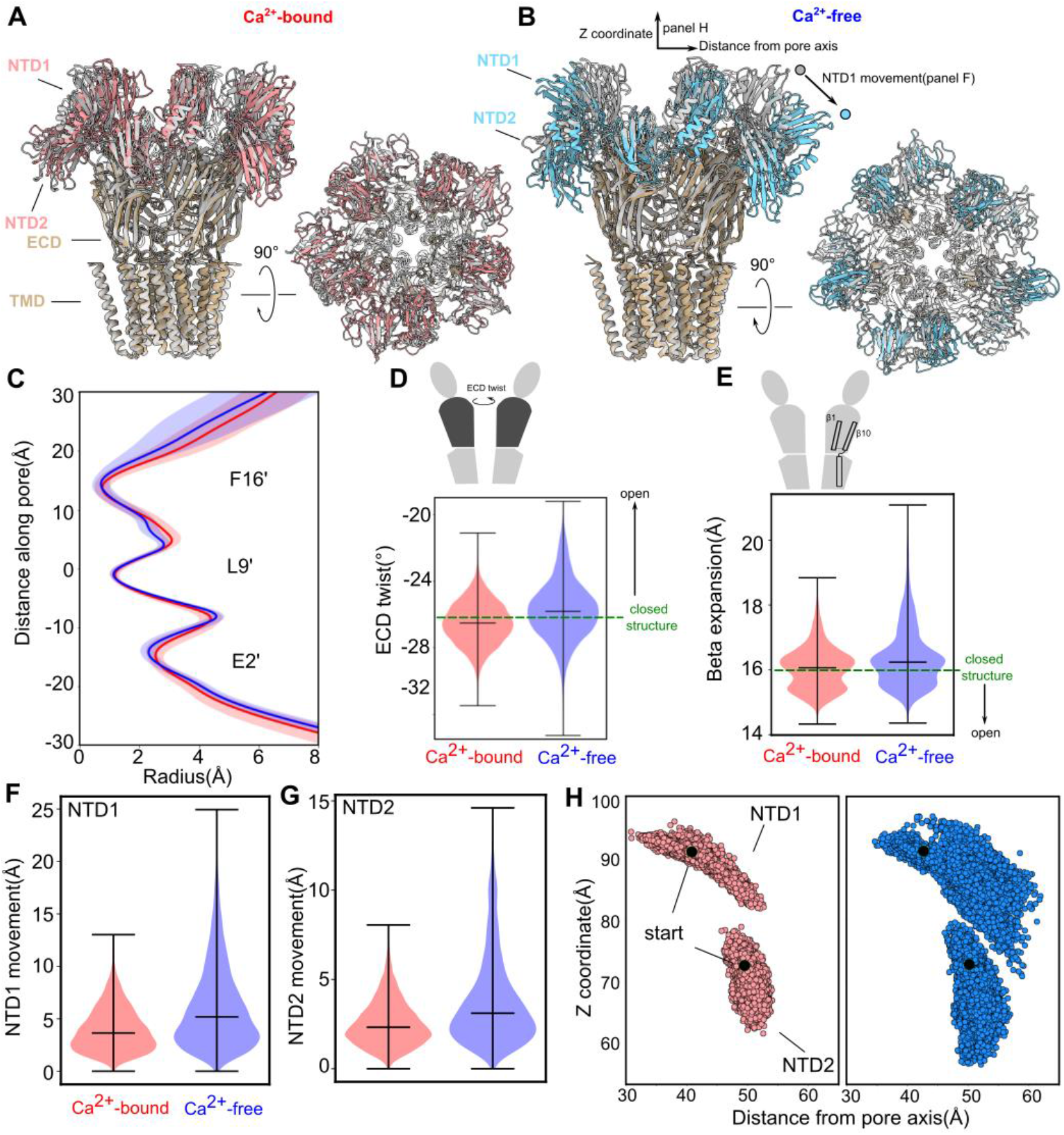
Destabilization of closed DeCLIC upon calcium depletion. **A)** Cryo-EM structure of DeCLIC in the presence of calcium (gray), superimposed with a snapshot from the end of an MD simulation of the same structure (tan), viewed from the membrane plane (left) and from the periplasm (right). For clarity, the NTD in the MD-snapshot is colored separately (red). **B)** Structure of DeCLIC in the sym calcium-free conformation (gray), superimposed with a snapshot from the end of an MD simulation of the same structure (tan), viewed as in A. The NTD in the MD-snapshot is colored separately (blue). Arrows indicate metrics used to track motions of individual NTD1/NTD2 lobes. **C)** Radius profiles of the TMD pore during MD simulations in the presence of calcium (red) and in the sym state in the absence of calcium (blue). Solid lines represent mean values, shaded areas represent standard error of the mean. **D)** Raincloud plots for ECD twist, measured as described in Methods and depicted as in the cartoon above. Dashed green line indicates value for the starting (closed) structures, with black arrow indicating anticipated transition towards opening. **E)** Raincloud plots for β expansion, measured as described in Methods and depicted as in the cartoon above. **F)** Raincloud plots of NTD1 COM RMSD, measured for each self-aligned subunit throughout four replicate trajectories in the presence (left, red) and absence (right, blue) of calcium. Horizontal lines indicate median values and extrema. **G)** Raincloud plots of NTD2 COM RMSD, measured as in F. **H)** COM position of each NTD lobe over time across all subunits and trajectories, showing z-coordinates as a function of distance from the pore axis, with receptors aligned on the pentameric TMD. The starting positions were highlighted by black beads. Both NTD1 (above) and NTD2 (below) sample a wider range of positions in the absence (right, blue) than in the presence of calcium (left, red).

### Structural homology of DeCLIC NTD to other prokaryotic regulatory domains

**Initial characterization of the DeCLIC structure identified mammalian, bacterial, and viral protein domains with jelly-roll folds comparable to individual NTD lobes**.^**9**^ To further explore NTD conservation in the context of Ca^2+^ interactions, we used the Dali server^15^ to identify similar structures in the Protein Data Bank. **Using our structures of the largely homologous** NTD1 and NTD2 lobes (Fig. 5A) as search models, **highest-ranking hits (Z-score > 9) included** regulatory domains from a variety of bacterial proteins, including the subtilisin-like serine protease Tk-SP from *Thermococcus kodakaraensis* (PDB ID 3AFG)^,16^ collagen-binding domain of *Clostridium histolyticum collagenase* (PDB ID 1NQD),^17^ and periplasmic lysozyme inhibitor PliG from *Aeromonas hydrophila* (PDB ID 4DZG)^18^ (Fig. 5B–D). Interestingly, relevant domains of Tk-SP and *Clostridium collagenase* each contain two calcium binding sites, with one site in Tk-SP directly overlapping Site 1 in DeCLIC NTD1 (Fig. 5B–C). Thus, the DeCLIC NTD appears to leverage an architecture common to prokaryotic regulatory domains, particularly calcium-binding regions.

**Figure 5.**
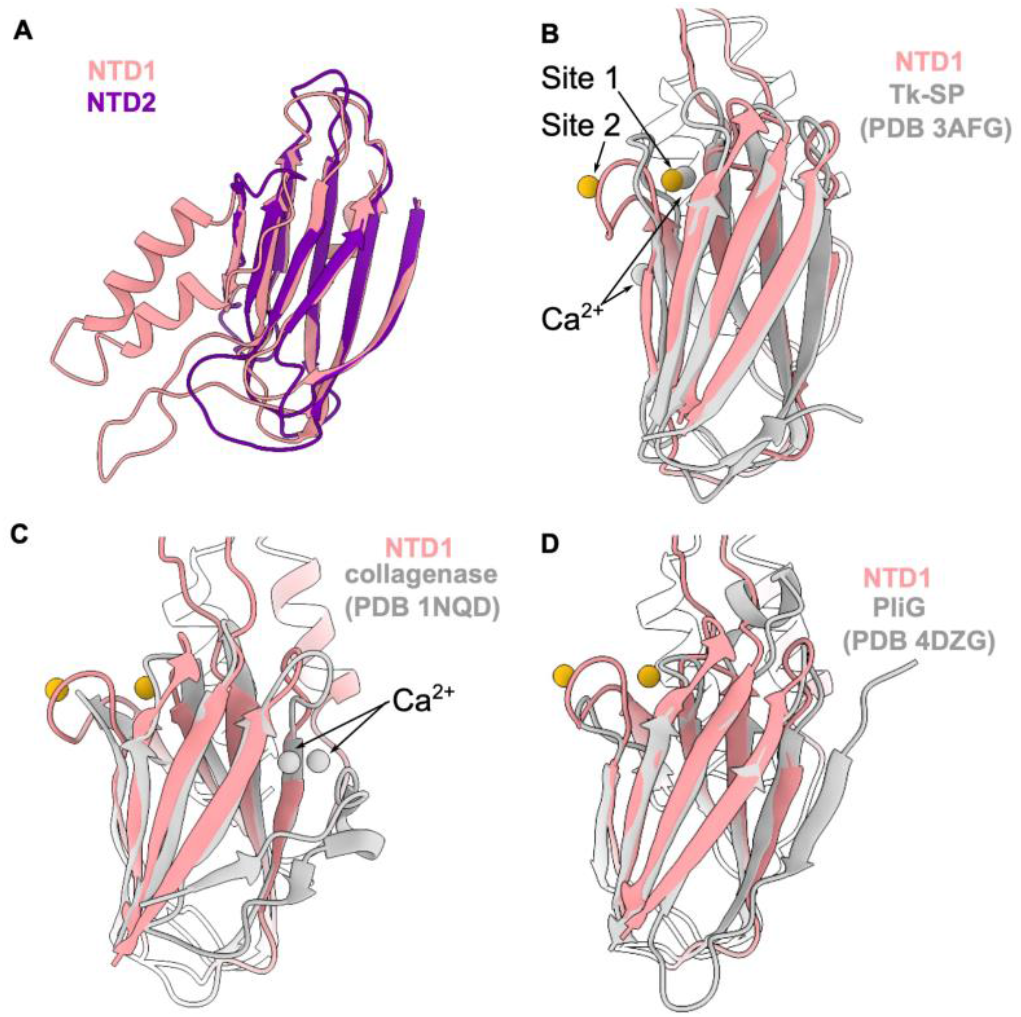
Structural homology of DeCLIC NTD to other prokaryotic regulatory domains. **A)** Superimposed structures of NTD1 (pink) and NTD2 (purple) without calcium, showing common β-sheet architecture. **B)** DeCLIC NTD1 (pink) with non-protein densities (gold), superimposed with Tk-SP with calcium (gray), showing similar Ca^2+^ binding at Site 1. **C)** DeCLIC NTD1 (pink) superimposed with the collagen-binding domain of *Clostridium collagenase* with calcium (gray). **D)** DeCLIC NTD1 (pink) superimposed with PliG (gray).

## Discussion

Differential dynamics underlie functional properties of many macromolecules, yet it remains challenging to characterize flexible or heterogeneous regions by classical structural methods.^12^ Here we modified sample preparation and data processing procedures to better resolve cryo-EM structures of a pLGIC under multiple conditions, including Ca^2+^ binding sites and a new asymmetrical conformation in the channel’s regulatory NTD. Specifically, we determined structures in lipid nanodiscs rather than detergent, and applied recent refinement tools including 3DFlex^12^ to incorporate neural networks in improving density maps representing flexible states.

Based in part on recent evidence from other bacterial pLGICs,^19^ one motivation for this work was to test whether reconstitution in lipid nanodiscs might reveal alternative functional states. Indeed, our structures include an asymmetric class in the absence of calcium, which provides direct structural evidence for transitions previously predicted by MD simulations and small-angle scattering.^10^ On the other hand, our nanodisc structures largely preserved the closed-pore TMD conformation in both the presence and absence of calcium. It is possible that differences in e.g. lipid composition between asolectin, *Xenopus* oocytes, *Escherichia* optimized for membrane protein expression, and native *Desulfofustis* membranes fundamentally alter the conformational landscape of DeCLIC, even in the context of relatively rigid circularized nanodiscs.^11^

Although the physiological relevance of NTD flexibility remains unclear, our data confirm and describe more fully Ca^2+^ binding at NTD as well as ECD subunit interfaces, one or both of which appear to stabilize the closed-pore structure particularly in the NTD. Structural and MD data supported relatively confident assignment of Ca^2+^ ions in the NTD (Site 1) as well as ECD (Site 3); additional non-protein densities observed in both domains (Sites 2, 4) may correspond to sites occupied at relatively low affinity and/or by alternative ligands, or in which the coordination is incompletely modeled by the present data and force fields. In particular, the highly negative potential of the DeCLIC surface may further promote accumulation of Ca^2+^ ions or other positively charged binding partners. In future work, it may be interesting to explore how calcium interactions or other electrostatic changes, for example protonation of acidic moieties by decreasing external pH, on the periplasmic surface of DeCLIC may influence channel structure and function.

More broadly, the NTD present in several prokaryotic pLGICs^8^ offers a promising model system to study regulatory protein domains. Individual NTD1/NTD2 lobes are structurally homologous to regulatory regions from a variety of prokaryotic proteins, including calcium binding domains in *Thermococcus* Tk-SP and *Clostridium* collagenase. N-terminal domains related to periplasmic binding proteins are found in eukaryotic ligand-gated channels including AMPA and NMDA receptors, where they are critical for agonist binding and allosteric regulation.^20,21^ Among eukaryotic pLGICs, the DeCLIC NTD offers a conceptual parallel to the regulatory ICD, for which no complete structures have yet been reported, possibly due to intrinsic disorder or the lack of an obligatory binding partner.^1^ It is plausible that the DeCLIC NTD is intrinsically dynamic in a physiological setting, or stabilized by interactions in the periplasm. Taken together, our results elucidate the structural basis for calcium binding and stabilization in a relatively dynamic pLGIC modulatory domain, with potential implications for allosteric and environmental regulation in a range of macromolecular systems.

## Methods

### Protein expression and purification

Expression and purification of DeCLIC was carried out as in previous publications.^10^ Briefly, the plasmid pET-20b containing the coding sequence of DeCLIC-MBP was transformed to C43(DE3) *Escherichia coli*, cells were were cultured overnight at 37 °C, then inoculated 1:100 into 2xYT media with 100 μg/mL ampicillin, and shaken until OD600 = 0.8 at 37 °C. After that, 100 μM isopropyl-β-D-1-thiogalactopyranoside (IPTG) was added to induce protein expression at 20 °C overnight.

Cells were harvested the next day and immediately resuspended in buffer A (20 mM Tris-HCl pH 7.4, 300 mM NaCl) supplemented with 1 mg/mL lysozyme, 20 μg/mL DNase I and protease inhibitors, then lysed by sonication on ice. Membranes were collected after ultracentrifugation and solubilized in buffer A with 2% n-dodecyl-β-D-maltoside (DDM) for 2 h at 4 °C. The solubilized membranes were ultracentrifuged, and supernatant incubated with amylose resin (NEB) with gentle rotation for 3 h at 4 °C. The amylose resin was loaded into a gravity column and washed in buffer A with 0.1% DDM. DeCLIC-MBP was eluted in buffer A with 0.02% DDM and 20 mM maltose, then further purified by gel-filtration chromatography in buffer A with 0.02% DDM. The peak fractions were collected and digested by thrombin to remove the MBP tag in 4 °C overnight. The digestion product was further purified by gel-filtration chromatography in buffer A with 0.02% DDM, collecting peak fractions corresponding to the predicted size of pentameric DeCLIC.

### Protein-nanodisc reconstitution

A plasmid containing the circularized nanodisc membrane scaffold protein spMSP1D1 was obtained from Addgene (#173482). Purification of spMSP1D1 followed previously published protocols.^11^ For reconstitution in circularized nanodiscs, purified DeCLIC, spMSP1D1 and soybean polar lipid extract (Avanti) were mixed at a 1:2:40 molar ratio, then incubated on ice for 1 h. Bio-Beads SM-2 resin (Bio-Rad) was added to the mixture, then gently rotated overnight at 4 °C. On the next day, the supernatant was collected and further purified by gel-filtration chromatography on a Superose 6 column in 20 mM HEPES pH 7, 150 mM NaCl. Peak fractions were pooled and concentrated to ∼5 mg/mL.

### Cryo-EM sample preparation and data collection

The nanodisc sample was mixed with stock solutions to reach a final concentration of 3 mM fluorinated Fos-choline-8 (Fos8-F, Anatrace) with 10 mM CaCl_2_ or 10 mM EDTA. For each dataset, 3 μL of the mixture was applied to a glow-discharged grid (R1.2/1.3 300 mesh Au grid, Quantifoil), blotted for 2 s with force 0, and plunged into liquid ethane using a Vitrobot Mark IV (Thermo Fisher Scientific).

Cryo-EM data were collected on a 300kV Titan Krios (Thermo Fisher Scientific) electron microscope with a K3 Summit detector (Gatan) at 105,000x magnification, corresponding to 0.8464 Å/px using the software EPU (Thermo Fisher Scientific). The total dose was ∼45 e^-^/Å^2^ and the defocus range −0.8 to −2.4 μm.

### Cryo-EM data processing

Dose-fractionated images in super-resolution mode were internally gain-normalized and binned by 2 in EPU during data collection. The initial data processing was done in Relion 3.0.7,^22^ including motion correction, contrast transfer function (CTF) estimation with CTFFIND 4.1, template particle picking, particle extraction, 2D classification, 3D classification, 3D refinement, CTF refinement and polishing. Two rounds of 2D classification were done to remove junk particles, and 3D classification (classes=4) was used to analyze structural heterogeneity. Particles from different classes were refined separately with symmetry C5 (with calcium, without calcium: *sym*) or C1 (without calcium: *asym*). Multiple rounds of CtfRefine and two of polishing were executed to improve resolution.

To improve density in the NTD, we continued the data processing in CryoSPARC v4.2.1.^23^ The particles after polishing were imported into CryoSPARC, and non-uniform refinement was run without imposing symmetry.^24^ 3DFlex refinement was applied with default settings including the training (laten dimensions 2) and flex reconstruct steps.^12^ The densities were further processed using DeepEMhancer 0.14 (mode tightTarget).^13^

### Model building and refinement

Model building was initiated by rigid-body fitting X-ray structures of DeCLIC (PDB ID 6V4S) and NTD1 (PDB ID 6V4B) into the densities. The models were manually checked and adjusted in Coot 0.9.5,^25^ and Ca^2+^, waters, and lipids were manually added. The resulting model was further optimized using real-space refinement in PHENIX 1.18.2 ^26^ and validated by MolProbity^27^.

Pore radius profiles were calculated using CHAP 0.9.1.^28^ Electrostatic potential was calculated by the “coulombic” function in UCSF ChimeraX. Structure figures were prepared using UCSF ChimeraX 1.3.^29^

### MD simulations

A summary of the MD simulation setup can be found in Table 2. DeCLIC nanodisc structures with calcium and without calcium: *sym* were embedded in bilayers of 680 1-palmitoyl-2-oleoyl-glycero-3-phosphocholine (POPC) lipids in CHARMM-GUI.^30^ The systems were subsequently solvated with TIP3P water and 150 mM CaCl_2_ (for the system with Ca^2+^) or 150 mM NaCl (for the system without Ca^2+^). The CHARMM36m force field was used to describe the proteins,^31^ with a multisite calcium model applied to the system with Ca^2+^.^32^

Simulations were performed using GROMACS 2020.5 ^33^ at 300 K using the velocity-rescaling thermostat^34^ and Parrinello-Rahman barostat^35^. The LINCS algorithm was used to constrain hydrogen-bond lengths,^36^ and the particle mesh Ewald method was used to calculate long-range electrostatic interactions.^37^ The systems were energy minimized and then equilibrated for 20 ns, with position restraints on the protein gradually released. Four replicates each of 300–400 ns were simulated as final unrestrained production runs.

Prior to analysis, MD simulation trajectories were aligned on the Cα atoms of the TMD region (residues 328-639) using MDAnalysis.^38^ Movement of the NTD domains (residues 35-195) was calculated by the COM movement of the Cα atoms in VMD^39^ and visualized with Matplotlib^40^. Measurements of ECD twist and β-expansion were adapted from previous studies of the prokaryotic pLGIC GLIC^14^, with ECD twist for each subunit calculated as the dihedral angle between a) the Cα COM of the ECD (residues 328–513) of one subunit, b) the equivalent ECD residues of all subunits, c) the TMD (residues 515–636) including all subunits, and d) the equivalent TMD residues of one subunit; β-expansion was calculated for each subunit as the COM distance between regions of β1 (residues 340–345) and β10 (residues 511–515).

## Data Availability

The cryo-EM maps and the corresponding atomic coordinates have been deposited in the Electron Microscopy Data Bank (EMDB) and the Protein Data Bank (PDB) under accession codes below. Non-uniform refinement: with Ca^2+^ (EMD-19994, PDB-9EV8), without Ca^2+^ *sym* (EMD-19997, PDB-9EVB), without Ca^2+^ *asym* (EMD-19995, PDB-9EV9). 3D-Flex refinement: with Ca^2+^ (EMD-19996, PDB-9EVA), without Ca^2+^ *sym* (EMD-19991, PDB-9EV1), without Ca^2+^ *asym* (EMD-19993, PDB-9EV7).

MD simulation parameters and trajectories are available at https://zenodo.org/records/10887632.

## Acknowledgements

We are deeply grateful to Professor Marc Delarue for inspiration and detailed feedback on this work. We thank Dr. Stavros Azinas for discussion of 3DFlex setup, members of Molecular Biophysics Stockholm for feedback on the project, and staff at the Cryo-EM Swedish National Facility for data collection and support. Cryo-EM data were collected at the Swedish National Facility funded by the Knut and Alice Wallenberg, Family Erling Persson and Kempe Foundations, SciLifeLab and Stockholm University. MD simulations were performed using the computing facilities of the Swedish National Infrastructure for Computing (SNIC 2022/3-40), and supported by BioExcel (EuroHPC grant no. 101093290). C.F. was supported by grant FV-5.1.2-0523-19 from Stockholm University, and R.J.H. and E.L. by grants from the Swedish Research Council (2019-02433, 2021-05806) and Swedish e-Science Research Center.

## Author contributions

C.F. performed the sample preparation, cryo-EM data collection and data, structural analysis and MD simulations. M. L. and Y.Z. helped with the MD simulations setup and analysis. R.J.H. and E.L. supervised the project. C.F., R.J.H. and E.L. contributed to the manuscript writing and revision.

