## Supplemental Data 1 for "Calcium stabilization of a flexible N-terminal domain in a pentameric ligand-gated ion channel"

### Supplemental Figures

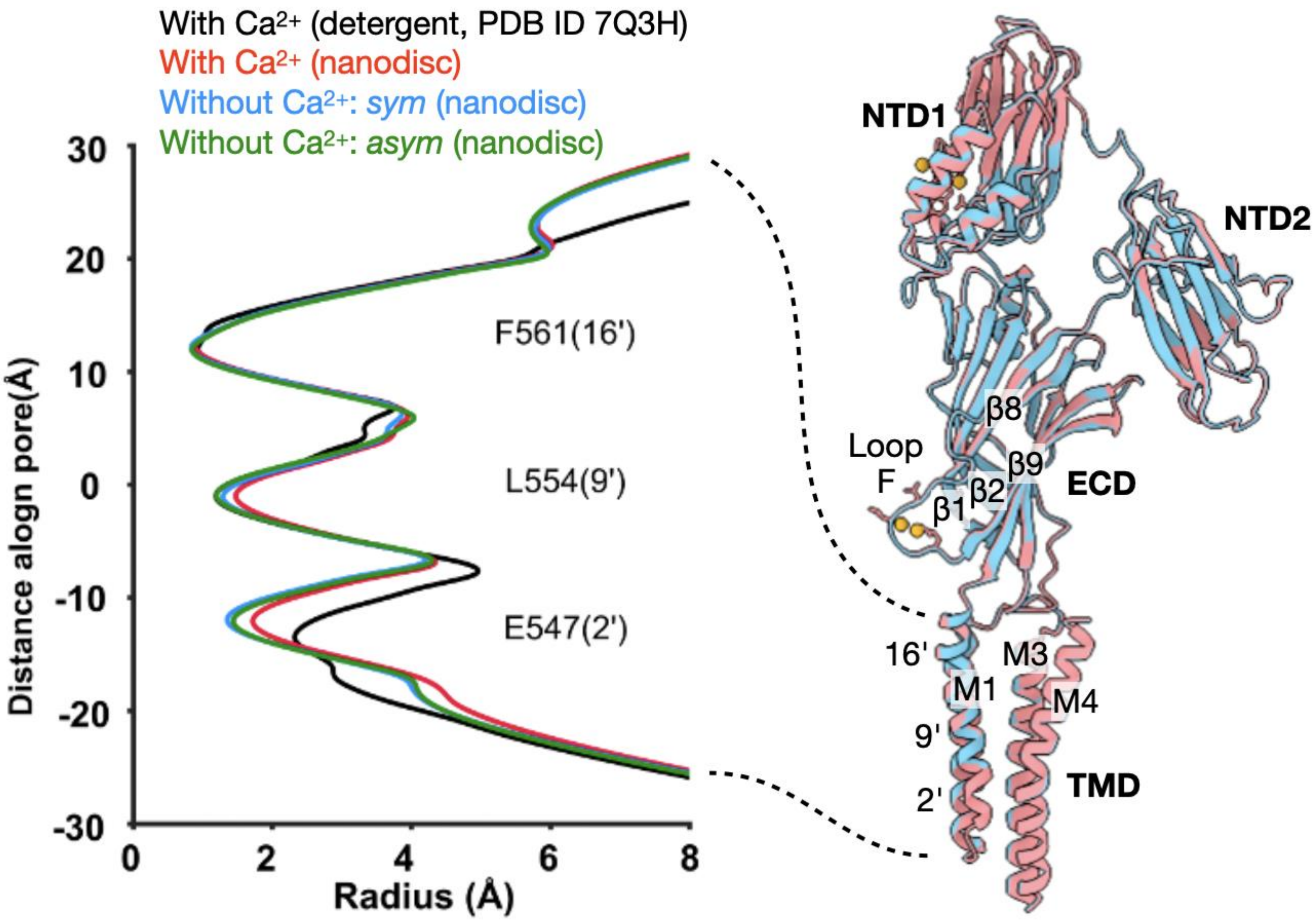

**Figure S1. Comparison of DeCLIC structures in the presence and absence of calcium.** *Left,* Radius profiles of the TMD pore in nanodisc structures in the presence of calcium (red) and in the *sym* (blue) and *asym* (green) states in the absence of calcium, as well as our previously reported $Ca^{2+}$-bound cryo-EM structure in detergent (black). *Right,* Superimposed structures of DeCLIC in the presence of calcium (red) and in the *sym* state in the absence of calcium (blue). For simplicity, a single subunit is shown, viewed from the membrane plane with key domains labeled. In the leftmost TMD helix (M2), key pore-facing residues at the 2', 9', and 16' positions are labeled.

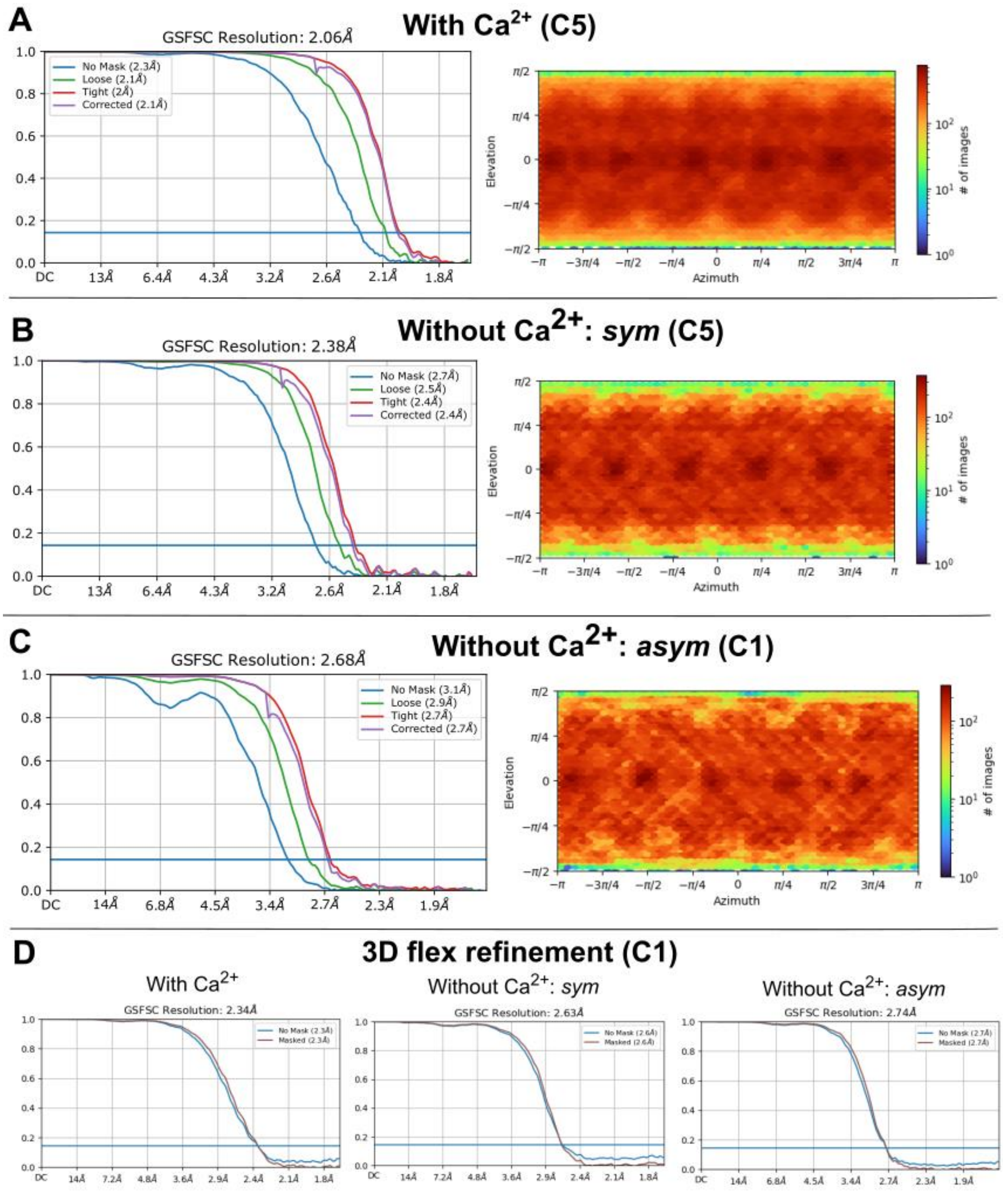

**Figure S2. Refinement plots for DeCLIC cryo-EM densities. A)** *Left,* Fourier shell correlations (FSC) versus overall resolution for DeCLIC in the presence of calcium, including correlations for the unmasked map (blue) and with a relatively loose (green), tight (magenta), or corrected (purple) mask applied in CryoSPARC. Solid blue line represents the gold-standard FSC (GSFSC, 0.143) indicator of true map resolution. *Right,* angular distribution of DeCLIC particles in the presence of calcium, colored according to scale bar at right. **B)** Plots as in *A* corresponding to the *sym* state in the absence of calcium, refined with C5 symmetry. **C)** Plots as in *A* corresponding to the *asym* state in the absence of calcium, refined without imposing symmetry. **D)** FSC of the 3D flex refinement maps.

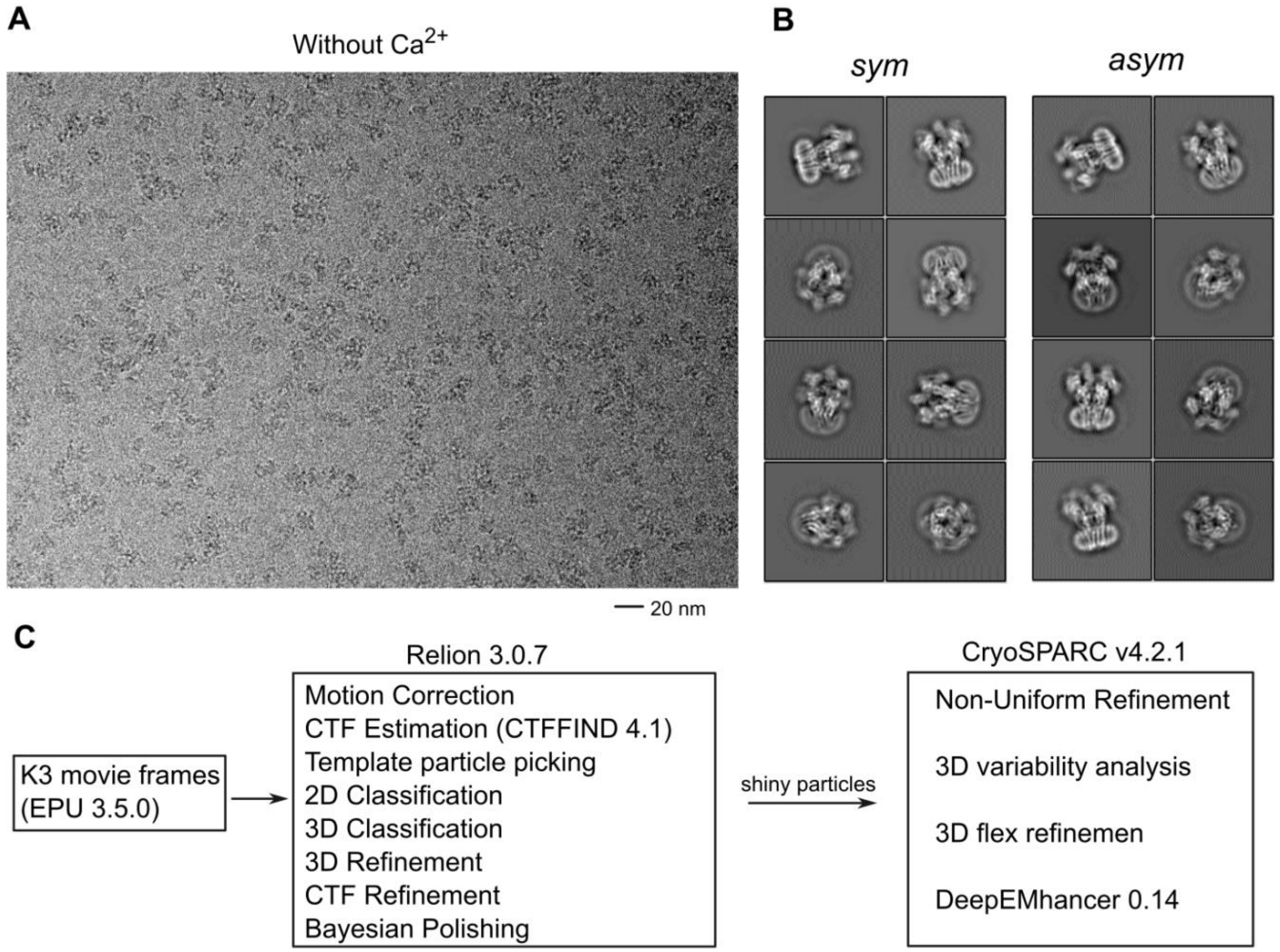

**Figure S3. Processing pipelines for DeCLIC structures. A)** Representative cryo-EM micrograph. **B)** Representative 2D classification images for classes designated *sym* (left) and *asym* (right) in the absence of calcium. **C)** Data processing workflow for all datasets.

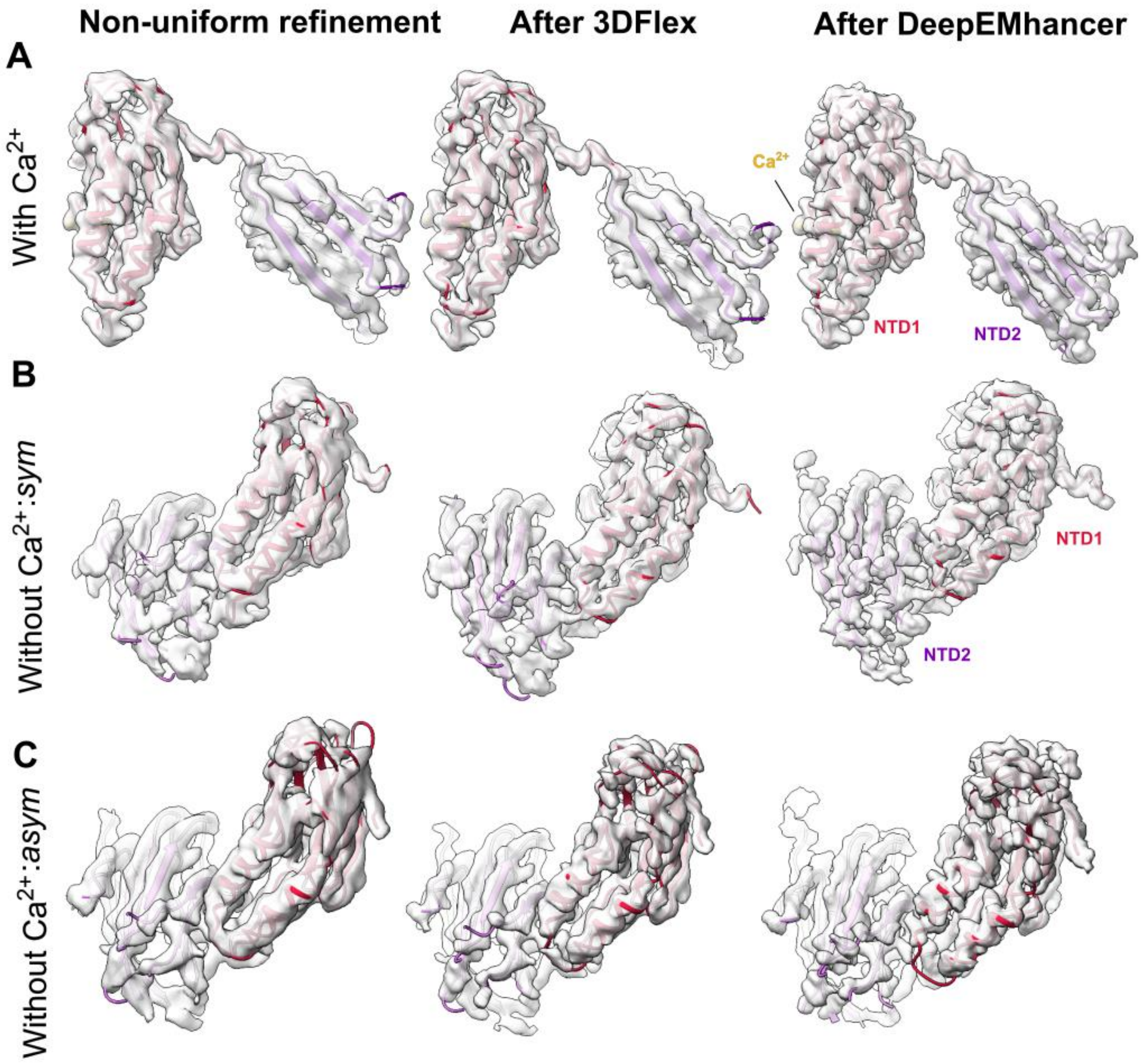

**Figure S4. Flexible refinement and deep learning-based post processing of NTD densities.** **A)** Cryo-EM maps associated with the NTD1–NTD2 region of a single DeCLIC subunit in the presence of calcium (left), showing improved definition after refinement using 3DFlex (center) and subsequent post-processing with DeepEMhancer (right). **B)** Cryo-EM maps of intersubunit NTD1-NTD2 interaction region for the *sym* state in the absence of calcium. View of NTD1 is equivalent to *A*, alongside NTD2 from the principal subunit in order to show an alternative relationship. **C)** Cryo-EM maps as in *B* for the *asym* state in the absence of calcium.

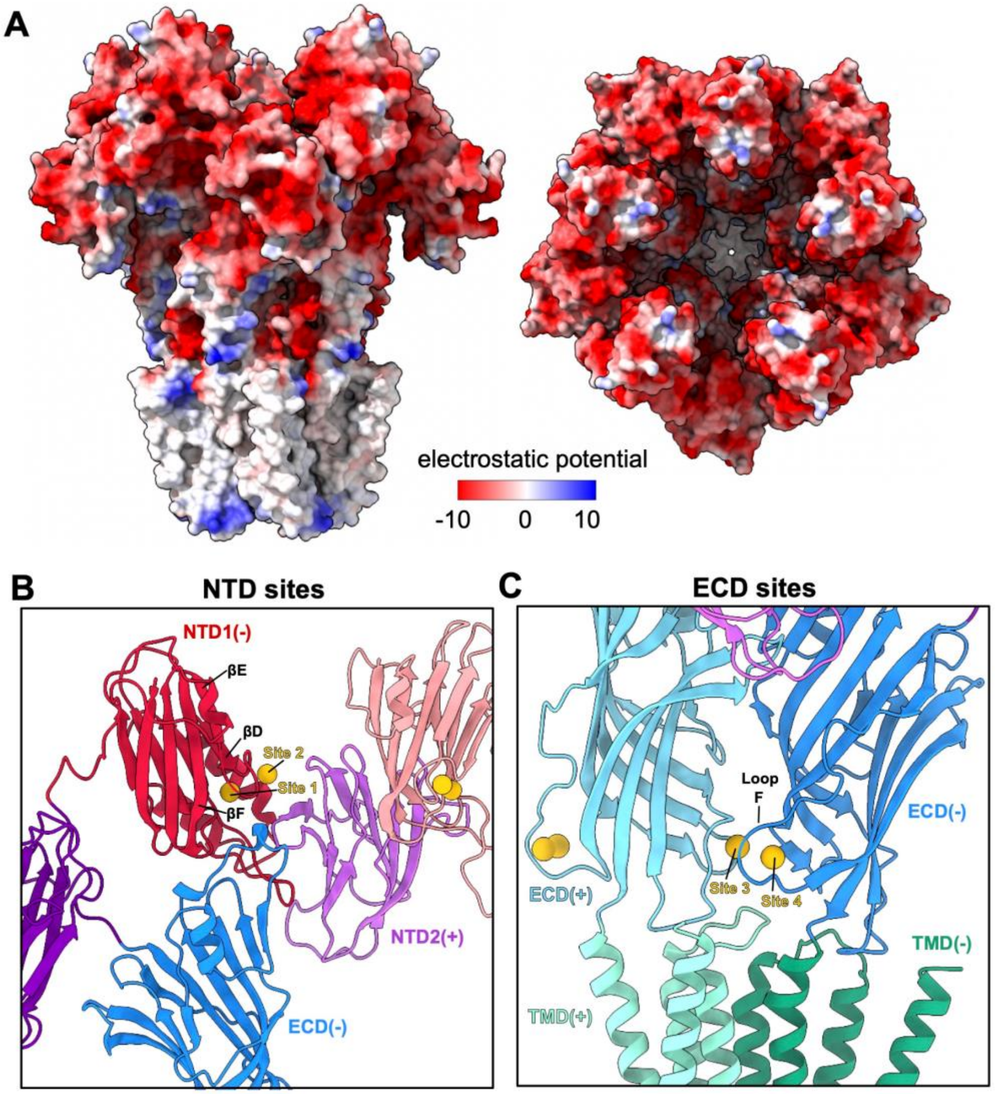

**Figure S5. Electrostatic potential and resolved calcium interactions in DeCLIC. A)** Surface representations of DeCLIC in the presence of calcium, viewed from the membrane plane (left) and from the periplasmic side (right). Molecular surfaces are colored by electrostatic potential according to center scale bar. **B)** Zoom view of an NTD $Ca^{2+}$ site, viewed from the periplasmic vestibule. For clarity, the complete complementary subunit is shown shaded darker (–, left), along with NTD lobes from the principal subunit shaded lighter (+, right). **C)** Zoom view of an ECD $Ca^{2+}$ site, viewed from the periplasmic medium. As in *B*, the principal subunit is shaded lighter (+, left), the complementary subunit darker (–, right). All models are colored as in *Fig. 1B.*

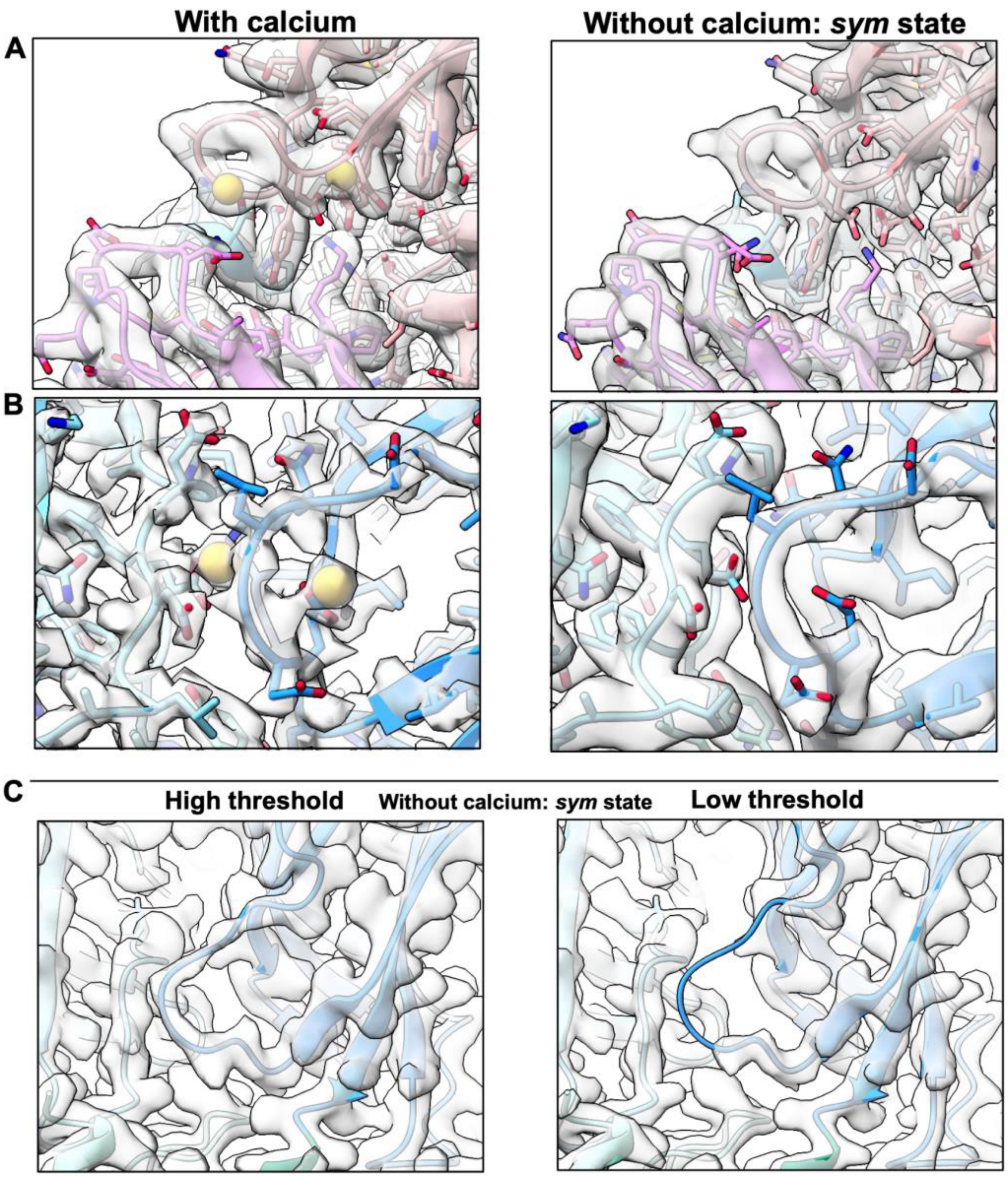

**Figure S6. Cryo-EM densities in calcium-binding regions of symmetric DeCLIC states in the presence and absence of calcium. A)** Cryo-EM maps surrounding a single NTD calcium site in the presence of calcium (left) and for the *sym* state in the absence of calcium (right), showing little structural change between conditions. **B)** Cryo-EM maps surrounding a single ECD calcium site in the presence of calcium (left) and for the *sym* state in the absence of calcium (right), showing a loss of definition particularly in loop F upon calcium dissociation. **C)** Cryo-EM maps as in *B* for the *sym* state in the absence of calcium, filtered at higher (left) and lower threshold (right) to reveal poor definition in loop F. All models are colored as in *Fig. 1B.*

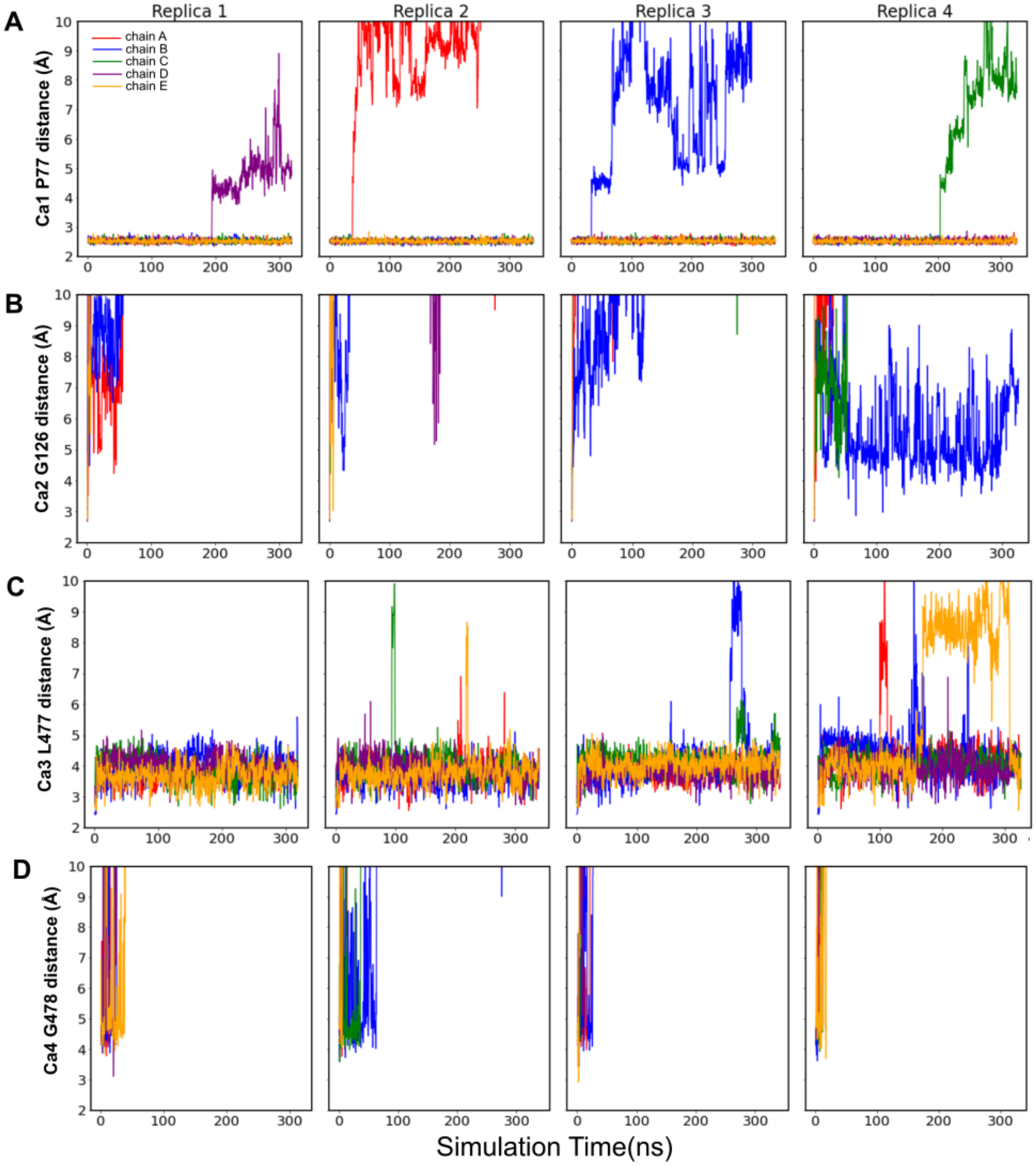

**Figure S7.** Dynamics **of calcium-binding in MD simulations.**
**A)** Dynamics of $Ca^{2+}$ binding in Site 1, quantified by the distance between the ion and the backbone carbonyl of P77 during four replicate MD simulations (left to right), each colored by chain. In each trajectory, $Ca^{2+}$ remained bound (within 3 Å of the coordinating atom) in four out of five chains. **B)** Dynamics of $Ca^{2+}$ binding in Site 2, quantified by the distance between the ion and the backbone carbonyl of G126, otherwise depicted as in *A*. All but one of the 20 $Ca^{2+}$ ions sampled dissociated >10 Å from the coordinating atom within 150 ns. **C)** Dynamics of $Ca^{2+}$ binding in Site 3, quantified by the distance between the ion and the backbone carbonyl of L477, otherwise depicted as in *A*. Despite transient displacements, $Ca^{2+}$ remained bound (within 5 Å of the coordinating atom) in all five chains at the end of all four trajectories. **D)** Dynamics of $Ca^{2+}$ binding in Site 4, quantified by the distance between the ion and the backbone carbonyl of G478, otherwise depicted as in *A*. All 20 $Ca^{2+}$ ions sampled dissociated >10 Å from the coordinating atom within 50 ns.

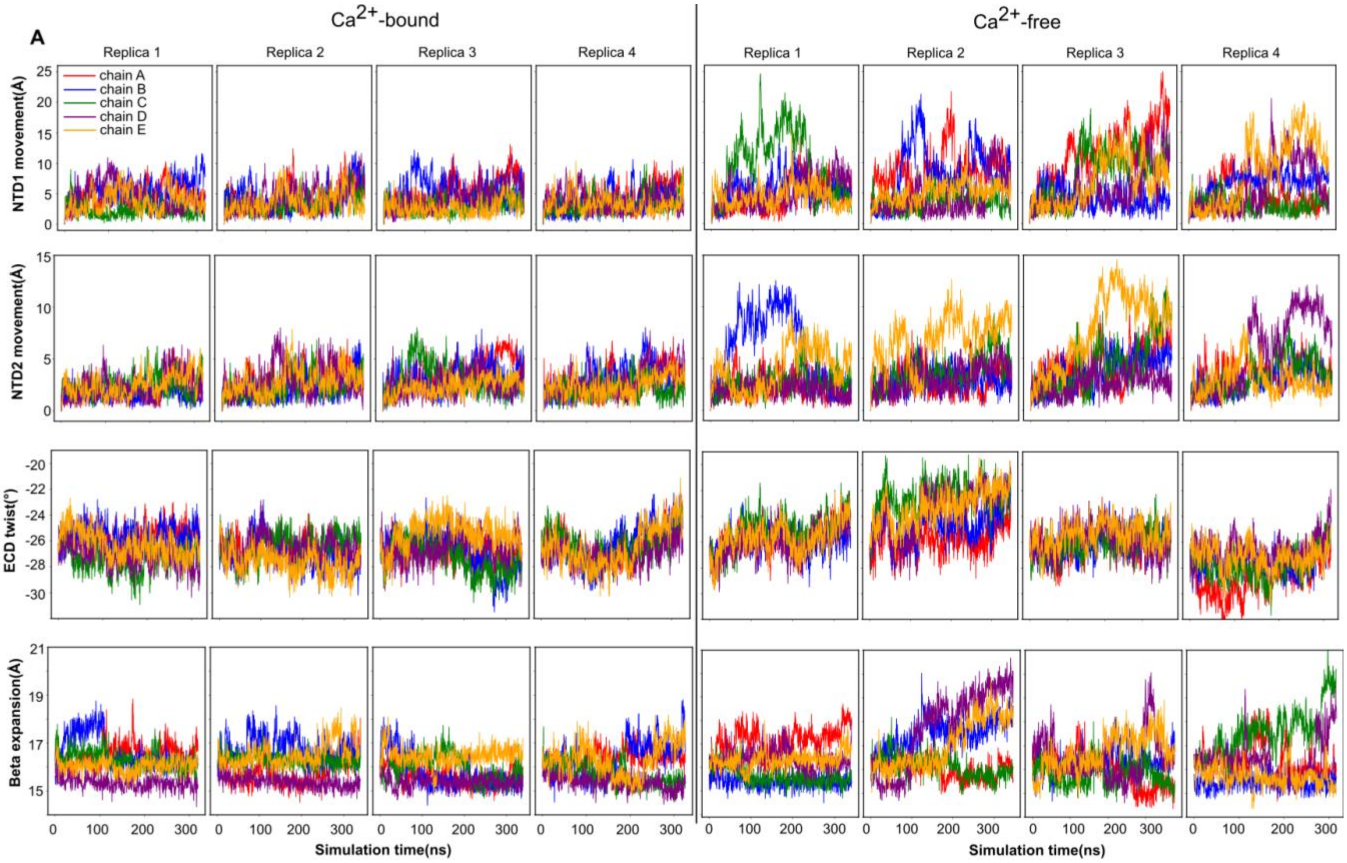

**Figure S8. Molecular dynamics of DeCLIC in the presence and absence of calcium. A)** Dynamics of each NTD1 lobe in four replicate simulations of DeCLIC, calculated by lobe center-of-mass RMSD and colored by chain, in the state determined with calcium (left) or in the *sym* state in the absence of calcium (right). **B)** Dynamics of each NTD2 lobe in the presence (left) and absence (right) of calcium, calculated and depicted as in *A*. **C)** ECD twist over time in the presence (left) and absence (right) of calcium, calculated as described in Methods and depicted as in *A*. **D)** β expansion over time in the presence (left) and absence (right) of calcium, calculated as described in Methods and depicted as in *A*. All measures show destabilization in the absence of calcium.

**Table 1 Cryo-EM data collection, refinement and validation statistics**

| | $Ca^{2+}$ | | EDTA *sym* | | EDTA *asym* | |
|---|---|---|---|---|---|---|
| | NU<br>(9EV8) | 3DFlex<br>(9EVA) | NU<br>(9EVB) | 3DFlex<br>(9EV1) | NU<br>(9EV9) | 3DFlex<br>(9EV7) |
| **Data collection and processing** | | | | | | |
| Magnification | 130,000 | 130,000 | 130,000 | 130,000 | 130,000 | 130,000 |
| Voltage (kV) | 300 | 300 | 300 | 300 | 300 | 300 |
| Electron exposure (e–/Å2) | 43.52 | 43.52 | 44.11 | 44.11 | 44.11 | 44.11 |
| Defocus range (μm) | -0.8 to -2.4 | -0.8 to -2.4 | -0.8 to -2.4 | -0.8 to -2.4 | -0.8 to -2.4 | -0.8 to -2.4 |
| Pixel size (Å) | 0.8464 | 0.8464 | 0.8464 | 0.8464 | 0.8464 | 0.8464 |
| Symmetry imposed | C5 | C1 | C5 | C1 | C1 | C1 |
| Final particles | 894,494 | 314,000 | 395,520 | 350,000 | 302,612 | 302,000 |
| Map resolution (Å)<br>FSC threshold | 2.06<br>0.143 | 2.34<br>0.143 | 2.38<br>0.143 | 2.63<br>0.143 | 2.68<br>0.143 | 2.74<br>0.143 |
| **Refinement** | | | | | | |
| Map sharpening *B* factor ($Å^2$) | -74.5 | - | -95.1 | - | -94.4 | - |
| Non-hydrogen atoms<br>Protein residues<br>Ligands | 24330<br>3005<br>55 | 23526<br>3005<br>20 | 24090<br>3005<br>35 | 23492<br>3005<br>0 | 22523<br>2840<br>35 | 22195<br>2840<br>0 |
| *B* factors ($Å^2$)<br>Protein<br>Ligand | 47.29<br>20.66 | 49.95<br>46.26 | 96.68<br>39.64 | 50.84<br>- | 51.00<br>39.64 | 51.00<br>- |
| R.m.s. deviations<br>Bond lengths (Å)<br>Bond angles (°) | 0.005<br>0.999 | 0.004<br>0.603 | 0.006<br>1.044 | 0.003<br>0.557 | 0.003<br>0.621 | 0.003<br>0.551 |
| Validation<br>MolProbity score<br>Clashscore<br>Poor rotamers (%) | 1.71<br>9.40<br>0.39 | 1.32<br>3.20<br>0.70 | 1.67<br>5.50<br>0.86 | 1.31<br>3.21<br>0.43 | 1.43<br>4.60<br>0.54 | 1.37<br>3.85<br>0.58 |
| Ramachandran plot<br>Favored (%)<br>Allowed (%)<br>Disallowed (%) | 96.66<br>3.34<br>0.00 | 96.73<br>3.07<br>0.20 | 95.99<br>3.67<br>0.33 | 96.76<br>3.01<br>0.23 | 96.82<br>3.00<br>0.18 | 96.82<br>3.00<br>0.18 |

**Table 2 System setup of MD simulations**

| | DeCLIC $CaCl_2$ | DeCLIC NaCl |
|---|---|---|
| Simulation box | 162Å x 162Å x 195Å | 162Å x 162Å x 195Å |
| Number of atoms | 478718 | 476272 |
| Number of waters | 112456 | 112524 |
| Number of ions | $Ca^{2+}$: 380, $Cl^-$: 610 | $Na^+$: 452, $Cl^-$: 302 |
| Number of lipids | 680 POPC | 679 POPC |
| Simulation time | 4 replicates, > 300 ns each | 4 replicates, > 300 ns each |
